# Assessment of the Effects of Muira Puama Root Extract on Biochemical Indices, and Testicular Histomorphology in Cyclophosphamide-treated Rats

**DOI:** 10.1101/2025.09.21.677620

**Authors:** Adesope Kingsley Adeyemi, Onaolapo Adejoke Yetunde, Onaolapo Olakunle James

**Affiliations:** Department of Pharmacology and Therapeutics, Ladoke Akintola University of Technology, Ogbomoso, Oyo State, Nigeria; Behavioural Neuroscience and Neurobiology Unit, Department of Anatomy, Ladoke Akintola University of Technology, Ogbomoso, Oyo State, Nigeria; Behavioural Neuroscience and Neuropharmacology Unit, Department of Pharmacology and Therapeutics, Ladoke Akintola University of Technology, Ogbomoso, Oyo State, Nigeria

**Keywords:** Gonadotoxicity, cyclophosphamide neurotoxicity, Muira puama

## Abstract

Cyclophosphamide (CYP), a commonly used anticancer drug, is limited in its therapeutic application by gonadotoxic effects driven by oxidative stress and inflammation. This study investigated the protective role of Muira puama (MP) extract against CYP-induced testicular damage in rats. Sixty rats were randomly assigned into six groups: Group A (control) received normal saline; Groups B and C were treated with MP (25 mg/kg and 50 mg/kg feed, respectively); Group D received CYP alone; and Groups E and F were co-administered CYP with MP (25 mg/kg and 50 mg/kg, respectively). Treatments lasted four weeks, after which body weight, feed intake, serum interleukin-10 (IL-10), Tumour necrosis factor-alpha (TNF-α), malondialdehyde (MDA), total antioxidant capacity (TAC), hormonal levels and testicular histology were evaluated. CYP administration (Group D) significantly reduced weight gain, feed intake, Testosterone and IL-10 levels, while elevating MDA and TAC (p < 0.05). Co-treatment with MP (Groups E and F) improved body weight, feed intake, and IL-10 levels, while reducing MDA and mitigating testicular histopathological damage. The higher MP dose (50 mg/kg) conferred greater protection. TAC values in MP co-treated groups were lower than with CYP alone, suggesting modulation of antioxidant responses. Histological analysis showed severe seminiferous tubule degeneration and basement membrane disruption in CYP-treated rats, whereas MP preserved testicular architecture, particularly at 50 mg/kg. In conclusion, Muira puama extract attenuates CYP-induced gonadotoxicity through antioxidant and anti-inflammatory mechanisms, supporting its potential as a protective agent against chemotherapy-related reproductive toxicity.

## 1. Introduction

Cyclophosphamide (CYP) is a potent chemotherapeutic and immunosuppressive agent extensively employed in the management of various cancers and autoimmune disorders due to its broad efficacy [1]. As an alkylating agent, CYP undergoes hepatic metabolism to yield active compounds such as acrolein and phosphamide mustard [2]. These metabolites exert cytotoxic effects by crosslinking DNA, inhibiting cell division, and triggering cell death. Additionally, they generate reactive oxygen species (ROS) and promote lipid peroxidation, processes that significantly contribute to tissue injury. In the male reproductive system, CYP exposure has been shown to induce testicular toxicity, largely through the accumulation of acrolein, which provokes oxidative stress, inflammation, and apoptosis [3]. The resulting damage manifests as impaired spermatogenesis, hormonal dysregulation, and reduced fertility, underscoring the need for protective interventions.

Recently, natural product–based therapeutic agents have attracted increasing attention as effective and safer alternatives for treating diseases and mitigating drug-induced toxicities [4]. Phytochemicals derived from medicinal plants possess diverse pharmacological properties, particularly antioxidant and anti-inflammatory activities, which render them valuable in conditions where oxidative stress plays a central role [5,6]. Such properties also highlight their promise as preventive or adjunctive agents in chemotherapy, where they may reduce oxidative damage and restore physiological balance.

Muira puama (MP), a medicinal plant with established antioxidant, anti-inflammatory, and adaptogenic properties, represents a potential candidate for ameliorating CYP-induced gonadotoxicity [7]. Evaluating its effects may not only offer insights into reproductive protection but also expand therapeutic strategies for managing chemotherapy-related adverse outcomes.

## 2. Methodology

### 2.1 Drugs and Reagents

Muira puama extract (Via Vittorio Venevento, 3/L25128 Brescia (BS) Italy), Cyclophosphamide injection (Endoxan-Asta®), Interleukin-10 assay kit (Biovision Inc., Milpitas, CA, USA). Tumour necrosis factor alpha (TNF α) assay kit (Biovision Inc., Milpitas, CA, USA). Total antioxidant capacity (TAC) assay kit (Biovision Inc., Milpitas, CA, USA).

### 2.2 Animals

The healthy Wistar rats utilised in this investigation were sourced from Empire Breeders, located in Osogbo, Osun State, Nigeria. Rats were housed in hardwood cages measuring 20 x 10 x 12 inches within room temperature (25°C ±2.5°C), with lights on at 7:00 am and off at 7 pm. Rats were granted unrestricted access to feed and water. All procedures were executed in compliance with the protocols of the Faculty of Basic Clinical Sciences, Ladoke Akintola University of Technology, adhering to the regulations for animal care and use outlined in the European Council Directive (EU2010/63) on scientific procedures involving living animals.

### 2.3 Experimental Methods

A total of Sixty rats weighing 120-150 g each were randomly assigned into six groups (n=10 each). Group A (control) received 10 ml/kg of normal saline, group B and C received 25 mg/kg and 50 mg/kg of Muira puama extract respectively, Group D (Cyclophosphamide) received 200 mg/kg of Cyclophosphamide, group E 200 mg/kg of CYP and 25 mg/kg of Muira puama extract, group F received 200 mg/kg of CYP and 50 mg/kg of Muira puama extract. CYP were administered intraperitoneally on days 1 and 2 while Muira puama extract was administered orally for 15 days. Doses of MP were determined based on evidences from previous research On day 16, rats were euthanized by cervical dislocation, and blood was drawn for analysis of biochemical parameters (TAC, MDA, IL-10 and TNF-α). The Testis were excised, fixed in 10% neutral buffered Bouin’s fluid, and testicular tissues sections were processed, embedded in paraffin, and stained for histological evaluation.

### 2.4 Assessment of body weight and feed intake

Body weight of animals in all groups was measured weekly using an electronic weighing scale (Mettler Toledo Type BD6000, Switzerland), while food consumption was assessed using a weighing balance as previously described-[9, 10]. The percentage change in body weight or food intake for each animal was calculated using the following equation, and the results for all animals were computed to find the statistical mean:

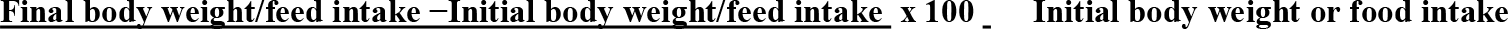

### 2.5 Biochemical Assays

#### 2.5.1 Measurement of malondialdehyde (MDA) level

Serum malondialdehyde (MDA) levels were determined following previously described protocols [11, 12]. Briefly, 200 µl of serum was incubated with 500 µl of thiobarbituric acid (TBA) reagent in a boiling water bath for 20 minutes. The reaction mixture was then cooled under running tap water for 13 minutes, followed by centrifugation at 3,000 rpm for 10 minutes to remove precipitates. The absorbance of the resulting supernatant was measured spectrophotometrically at 532 nm against a reagent blank (containing all reagents except serum), using a Shimadzu UV–VIS Recording 2401 PC®. The concentration of MDA was calculated using the formula: MDA (mol/ml) = (Absorbance of sample / Absorbance of standard) ×100

#### 2.5.2 Measurement of Total Antioxidant Status

Total antioxidant capacity was determined using the ferric reducing antioxidant power (FRAP) assay, as previously described [13, 14]. Briefly, 8 µl of serum was added to 240 µl of freshly prepared FRAP reagent and incubated at 37 °C for a few minutes. The absorbance of the reaction mixture was then measured at 532 nm against a reagent blank, using a Shimadzu UV–VIS Recording 2401 PC®. Results were expressed as mmol/L of Fe^2^□ equivalents.

#### 2.5.3 Measurement of Inflammatory Cytokines

Serum levels of tumour necrosis factor-α (TNF-α) and interleukin-10 (IL-10) were quantified using enzyme-linked immunosorbent assay (ELISA). Commercially available ELISA kits (Enzo Life Sciences Inc., NY, USA) were employed to determine the total concentrations (both bound and unbound forms) of each cytokine. The procedure was conducted according to established protocols [15,16].

### 2.6 Gonadosomatic Index (GSI)

Body weight (g) at the end of the experimental period and testes weight (g) taken at dissection were used to calculate GSI as previously described [17, 18]. GSI was determined by di-viding testicular weight by body weight.

### 2.7 Hormone Analysis

Blood was taken via a cardiac puncture and centrifuged immediately to separate plasma from cells. Total plasma testosterone was then measured using a commercially available radio-immunoassay (RIA) kit (Tianjin Medical & Bio-engineering Co., Ltd), as previously described [19]. The lower limit of detection for the kit was 0.04 ng/ml, while intra-and inter-assay variation coefficients were less than 5%. Luteinizing hormone and follicle stimulating hormone levels were measured using commercially available kits following the instructions of the manufacturer

### 2.8 Tissue Histology

Formalin-fixed samples of the testes were embedded in paraffin and sectioned at 5µm thickness. The sections were then dewaxed and rehydrated before being mounted on slides and stained with haematoxylin-eosin (H&E) for general histological study. The sections of the testes were examined microscopically using a Sellon-Olympus trinocular microscope (XSZ-107E, China) with a digital camera (Canon PowerShot 2500), and photomicrographs were taken. Histopathological examination was conducted by a technician blinded to the groupings.

### 2.9 Statistical Analysis

Data were analysed using Chris Roden s ez ANOVA for Windows with one-way ANOVA. One-way ANOVA was used to test the hypothesis, and Tukey’s HSD test was employed for post-hoc analysis. Results were expressed as mean ± S.E.M., and p<0.05 was considered significant.

## 3. Results

### 3.1 Effect of Muira puama root extract on body weight

Figure 3.1 shows the shows the effect of Muira puama extract on body weight measured as relative change in body weight in rats treated with cyclophosphamide. There was a significant (p < 0.001) increase in body weight with Muira puama (MP) at 25, and 50 and a decrease with CYP, and the CYP/MP25 compared to control. Compared to CYP, relative body weight increased with CYP/MP25 and CYP/MP50.

### 3.2 Effect of Muira Puama on feed intake

**Figure** shows the effect of Muira puama extract on relative change in feed intake in cyclophosphamide treated rats. There was a significant (p < 0.001) increase in feed intake with MP25and MP 50 and a significant decrease with CYP, CYP/MP25 and CYP/MP50 compared to control. Compared to CYP, feed intake increased with CYP/MP25 and CYP/MP50 respectively.

### 3.3 Effect of Muira puama on Biochemical parameters

#### 3.3.1 Effect of Muira Puama Extract on Oxidative stress parameters

**Table 1** shows the effect of Muira puama extract on serum Superoxide dismutase (SOD) activity, Glutathione peroxidase (GSH) activity, Total antioxidant capacity (TAC) and lipid peroxidation measured as malondialdehyde (MDA) Superoxide dismutase activity decreased significantly with MP25, MP50 and increased with CYP, CYP/MP25, and CYP/MP50, compared to control. Compared to CYP, SOD activity decreased significantly with CYP/MP25, and CYP/MP50. Glutathione peroxidase activity increased significantly with MP25, MP50 and decreased with CYP, CYP/MP25, and CYP/MP50, compared to control. Compared to CYP, GSH activity increased significantly with CYP/MP25, and CYP/MP50.

Total antioxidant capacity increased significantly with MP25, MP50, CYP/MP25, and CYP/MP50, and decreased with CYP compared to control. Compared to CYP, TAC levels increased significantly with CYP/MP25, and CYP/MP50. Malondialdehyde (MDA) levels increased significantly in groups administered MP 50 mg/kg and CYP compared to the control group. Malondialdehyde levels decreased with MP25 and MP50, and increased with CYP, CYP/MP25 and CYP/MP50 compared to control. Compared to CYP, MDA levels decreased significantly with CYP/MP25, and CYP/MP50.

#### 3.3.2 Effect of Muira Puama Extract on Inflammatory Cytokines and Caspase-3 levels

**Table 2** shows the effects of Muira puama on inflammatory cytokines and caspase-3 levels in cyclophosphamide-treated rats. Tumor Necrosis Factor (TNF-α) decreased significantly with MP25, MP50, CYP/MP25 and CYP/MP50 and increased significantly with CYP compared to control. Compared to CYP, TNF-α levels decreased significantly with CYP/MP25, and CYP/MP50.

Interleukin-10 levels increased significantly with MP25, MP50, CYP/MP25 and CYP/MP50 and decreased significantly with CYP compared to the control. Compared to CYP, IL-10 levels increased significantly with CYP/MP25 and CYP/MP50.

Caspase-3 levels decreased significantly with MP25, MP50 and increased with CYP, CYP/MP25 and CYP/MP50 compared to control. Compared to CYP, caspase-3 levels decreased significantly with CYP/MP25, and CYP/MP50.

#### 3.3.3 Effect of Muira Puama on hormonal levels

Plasma testosterone levels increased significantly (*p* < 0.001) with MP25, MP50, CYP/MP25 and CYP/MP50 and decreased with CYP compared to control. Compared to CYP, TAC levels increased significantly with CYP/MP25, and CYP/MP50.

Luteinizing hormone (LH) levels decreased significantly (*p* < 0.001) with CYP, CYP/MP25 and CYP/MP50 compared to control. Compared to CYP, LH levels increased with CYP/MP25 and CYP/MP50. Follicle stimulating hormone (FSH) levels decreased significantly (*p* < 0.001) with CYP, CYP/MP25 and CYP/MP50 compared to control. Compared to CYP, FSH levels increased with CYP/MP25 and CYP/MP50

**Table 1:** Effects of Muira Puama on oxidative stress parameters.

| Groups | TAC<br>(pg/ml) | MDA<br>(pg/ml) | SOD U/ml | GSH U/ml |
| --- | --- | --- | --- | --- |
| **Control** | 4.58±<br>0.04 | 4.75±<br>0.01 | 0.46±<br>0.01 | 4.75±<br>0.01 |
| **MP25** | 6.77±<br>0.03 | 2.76±<br>0.14* | 0.23±<br>0.01* | 2.76±<br>0.14* |
| **MP50** | 7.20±<br>0.11* | 1.28<br>±0.33* | 0.30±<br>0.02* | 1.28±<br>0.33* |
| **CYP** | 2.22±<br>0.01* | 22.6±<br>0.30* | 7.23±<br>0.25* | 22.6±<br>0.30* |
| **CYP/MP25** | 6.43±<br>0.09<sup>#</sup> | 8.58±<br>0.02<sup>#</sup> | 3.22±<br>0.09<sup>#</sup> | 8.58±<br>0.02<sup>#</sup> |
| **CYP/MP50** | 6.83±<br>0.06<sup>#</sup> | 7.24±<br>0.02<sup>#</sup> | 2.42±<br>0.26<sup>#</sup> | 78.24±<br>0.02<sup>#</sup> |

Data are represented as mean ± S.E.M., \* $p < 0.05$ significant difference from control. <sup>#</sup> $p < 0.005$ significant difference from CYP. Numbers of rats per treatment group=10. CYP: Cyclophosphamide, MP: Muira puama. SOD: Superoxide dismutase, GSH: Glutathione, Malondialdehyde, Total antioxidant capacity.

**Table 2:** inflammatory cytokines and Caspase -3 levels.

| Groups | TNF-α<br>(pg/ml) | IL-10<br>(pg/ml) | Caspase-3<br>(ng/ml) |
| --- | --- | --- | --- |
| Control | 23.16±0.34 | 51.42±6.52 | 0.52±0.002 |
| MP25 | 22.06±0.22<sup>*</sup> | 61.14±0.44<sup>*</sup> | 0.38±0.001<sup>*</sup> |
| MP50 | 22.15±0.41<sup>*</sup> | 74.87±1.61<sup>*</sup> | 0.26±0.001<sup>*</sup> |
| CYP | 40.20±0.65<sup>*</sup> | 18.54±1.80<sup>*</sup> | 3.65±1.00<sup>*</sup> |
| CYP/MP25 | 16.12±0.15<sup>*#</sup> | 61.73±1.88<sup>*#</sup> | 1.23±0.01<sup>*#</sup> |
| CYP/MP50 | 15.22±0.30<sup>*#</sup> | 71.01±4.64<sup>*#</sup> | 1.01±0.02<sup>*#</sup> |

Data are represented as mean ± S.E.M., \* $p$ <0.05 significant difference from control. # $p$ <0.005 significant difference from CYP. Numbers of rats per treatment group=10. CYP: Cyclophosphamide, MP: Muira puama. TNF-α : Tumour Necrosis Factor alpha IL: Interleukin-10 levels

### 3.4 Effect of Muira Puama on Gonadosomatic index

Figure 3 shows the effect of Muira puama extract on gonadosomatic index in CYP-induced gonadal toxicity in rats. Gonadosomatic index (GSI) decreased significantly (*p* < 0 .001) with CYP, CYP/MP25 and CYP/MP50 compared to control. Compared to CYP, GSI increased significantly with CYP/MP25 and CYP/MP50.

**Figure 1:**
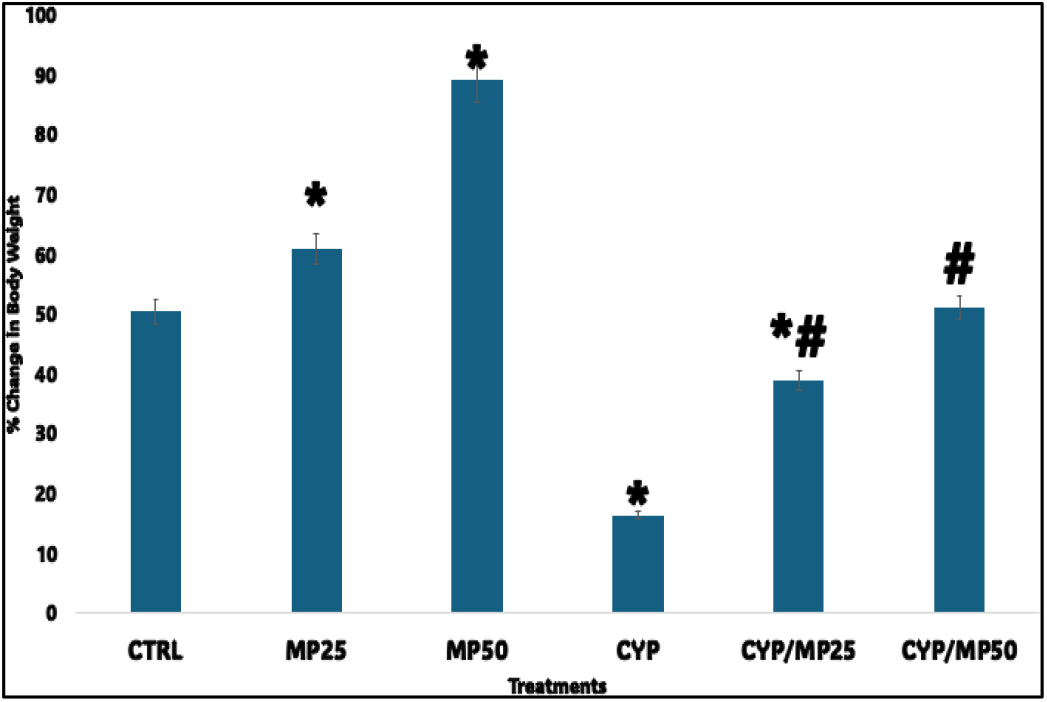
Effect of Muira puama on relative change in body weight in cyclophosphamide treated rats. Each bar represents Mean ± S.E.M, *p<0.05 significant difference from control. #p<0.05 significant difference from CYP. number of rats per treatment group =10. CYP: Cyclophosphamide. MP: Muira puama

**Figure 2:**
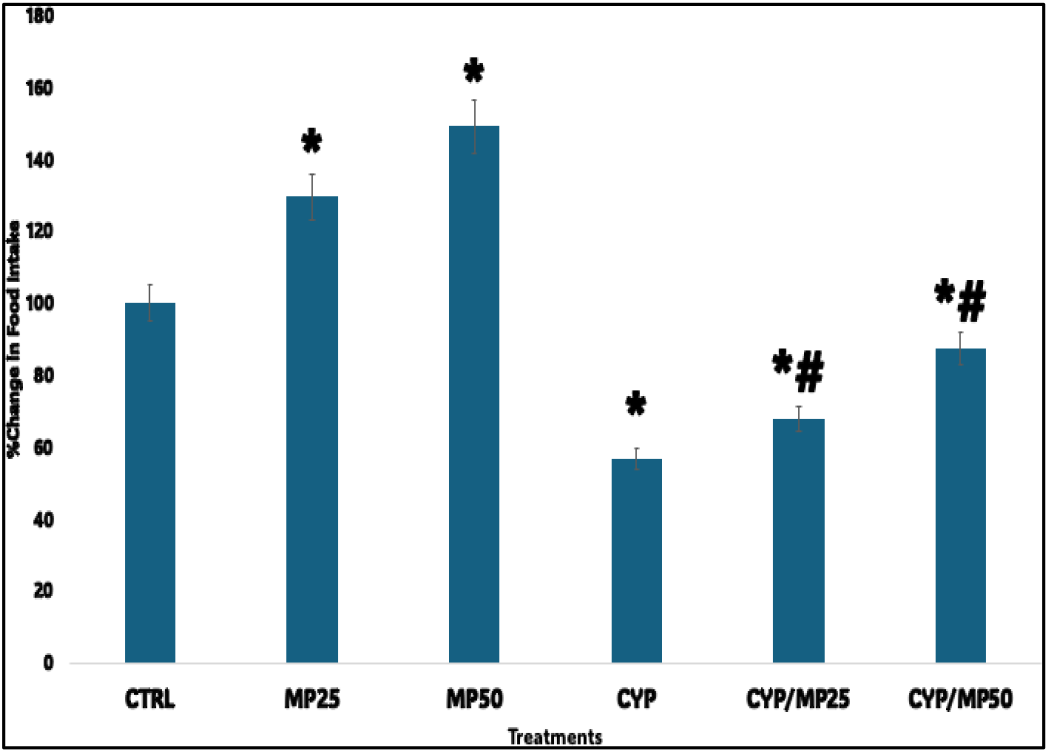
Effect of Muira puama extract on relative change in feed intake in cyclophosphamide treated rats. Each bar represents Mean ± S.E.M, *p<0.05 significant difference from control. #p<0.05 significant difference from CYP. Number of rats per treatment group =10. CYP: Cyclophosphamide. MP: Muira puama

**Figure 3:**
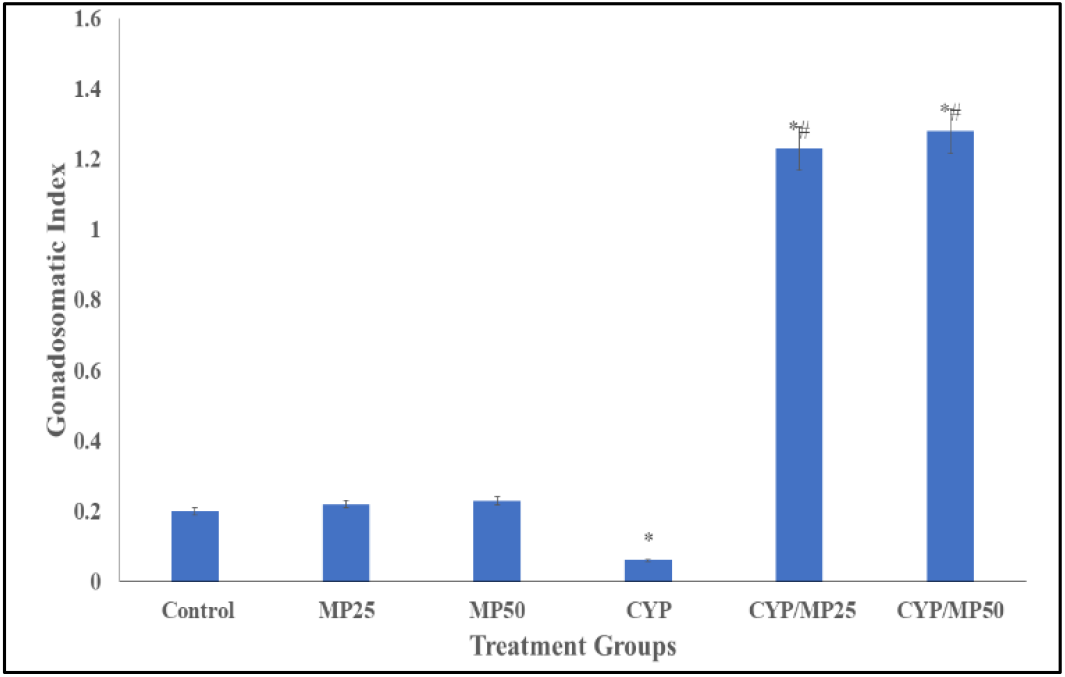
Effect of Muira puama extract on gonadosomatic index in cyclophosphamide treated rats. Each bar represents Mean ± S.E.M, *p<0.05 significant difference from control. #p<0.05 significant difference from CYP. Number of rats per treatment group =10. CYP: Cyclophosphamide. MP: Muira puama

### 3.5 Effect of Muira Puama on Testicular Histology

Plate 1(A-F) represents haematoxylin and eosin (H&E)-stained sections of the rat testes. Examination of the slides from the control group (A) revealed normal testicular histoarchitecture, characterized by well-defined Leydig cells, myoid cells, Sertoli cells, lumen, and a well-shaped basement membrane. These features are consistent with healthy and functional testes.

In contrast, the cyclophosphamide-treated group (D) exhibited histological alterations, including disorganized seminiferous tubules, reduced germ cell population, and a disruption of the basement membrane, indicating significant gonadotoxicity However, the groups treated with Muira puama extract (E and F) demonstrated varying degrees of histological preservation. Plates 1E and 1F showed better structural organization of the seminiferous tubules, improved Leydig cell integrity, and the presence of spermatogenic cells, indicating partial protection against cyclophosphamide-induced damage.

**Table 3:** Effect of Muira Puama on Hormone levels.

| Groups | T (ng/ml) | LH<br>(mIU/ml) | FSH (mIU/ml) |
| --- | --- | --- | --- |
| Control | 1.16±0.04 | 5.65±0.13 | 0.60±0.001 |
| MP25 | 2.02±0.02<sup>*</sup> | 5.30±0.12 | 0.54±0.001<sup>*</sup> |
| MP50 | 2.23±0.04<sup>*</sup> | 5.35±0.10 | 0.53±0.001<sup>*</sup> |
| CYP | 0.30±0.01<sup>*</sup> | 1.26±0.02<sup>*</sup> | 0.10±0.001<sup>*</sup> |
| CYP/MP25 | 2.80±0.15<sup>*#</sup> | 3.70±0.20<sup>*#</sup> | 1.30±0.001<sup>*#</sup> |
| CYP/MP50 | 2.87±0.12<sup>*#</sup> | 4.01±0.11<sup>*#</sup> | 1.46±0.002<sup>*#</sup> |

Data are represented as mean ± S.E.M., \* $p$ <0.05 significant difference from control. # $p$ <0.005 significant difference from CYP. Numbers of rats per treatment group=10. CYP: Cyclophosphamide, MP: Muira puama. T: Testosterone, FSH: Follicle Stimulating Hormone, LH: Luteinizing Hormone.

## 4. Discussion

In this study, cyclophosphamide administration was associated with a significant reduction in body weight relative to control, which is consistent with previous studies that had reported cytotoxic and cachexia-inducing effects of CYP ([17, 18, 21, 22]. Also in this group, a reduction in feed intake was observed consistent with the reports that had shown that CYP impairs appetite, alters gastrointestinal integrity, and increases oxidative stress, all of which have been shown to contribute to weight loss in experimental animals and cachexia in humans [23, 24]. In contrast, administration of Muira puama (MP) extract at 25 and 50 mg/kg significantly increased relative body weight and feed intake when given alone, suggesting a potential nutritive, anabolic, or appetite-stimulating effect. This aligns with ethnomedicinal reports of Muira puama as a tonic and adaptogen, capable of enhancing energy balance and general vitality [8]. Also, in CYP/MP co-treatment groups, relative body weight was significantly higher than with CYP alone, particularly in CYP/MP50. This implies that MP had the capacity to confer a protective effect against CYP-induced weight loss. Possible mechanisms may include its ability to reduce oxidative damage and preserving tissue integrity, improving feed intake suppressed by CYP which was also observed in this study. It could also have the capacity to enhance nutrient utilization and energy homeostasis. The partial restoration in the CYP/MP25 group, though not as strong as in CYP/MP50, suggests a dose-dependent effect. However, the fact that CYP/MP25 still showed lower weight compared to control indicates that the protective effect at lower doses may be insufficient to fully counteract CYP toxicity

Cancer chemotherapy has been reported to both promote and hinder treatment by inducing oxidative stress through the generation of reactive oxygen species (ROS). The generation of oxidative stress is considered one of the mechanisms by chemotherapies acts against cancerous cells, resulting in DNA damage and cell death. However, excessive oxidative stress can interfere with the effectiveness of chemotherapy and cause significant side effects [25]. In this study CYP) treatment resulted in a distinct oxidative stress profile characterized by increased SOD activity, reduced GSH activity, reduced TAC, and elevated MDA levels. These findings are consistent with the known pro-oxidant effects of CYP, which generates ROS and disrupts antioxidant enzyme balance, culminating in lipid peroxidation and membrane damage [21, 25] Administration of Muira puama (MP) extract alone exerted differential effects on antioxidant status. At both 25 and 50 mg/kg, Muira puama significantly increased GSH activity and enhanced TAC, indicating a stimulatory effect on endogenous antioxidant defense. The SOD activity was significantly reduced in the MP groups compared to control. This paradoxical decrease may reflect a reduced requirement for SOD-mediated dismutation of superoxide due to MP’s direct free radical scavenging ability or enhancement of downstream antioxidant systems. In co-treatment groups (CYP/MP25 and CYP/MP50), antioxidant balance was partially restored. Relative to CYP alone, SOD activity decreased toward control levels, while GSH activity and TAC were significantly elevated, indicating a corrective effect of MP against CYP-induced antioxidant suppression. The reduction in SOD activity relative to CYP may suggest normalization of enzyme activity rather than outright inhibition, reflecting lower oxidative burden due to MP’s intervention. Lipid peroxidation, measured as MDA levels, further highlights this protective effect. While CYP markedly increased MDA (consistent with heightened oxidative membrane damage), co-administration with MP (both CYP/MP25 and CYP/MP50) significantly reduced MDA, indicating attenuation of lipid peroxidation. The MP-alone groups also showed reduced MDA levels, particularly at 25 mg/kg, supporting a direct antioxidative and membrane-protective effect.

Cyclophosphamide has been shown to exert complex and at times opposing effects on inflammatory cytokines. On one hand, it has been associated with triggering pro-inflammatory responses by inducing cell death. However, at lower doses it may exert anti-inflammatory actions by suppressing distinct T helper responses and promoting populations that release anti-inflammatory cytokines such as IL-10 and transforming growth factor-β [26]. This dual response often described as the “Janus face” of cyclophosphamide [27]. In this study, CYP’s ability to trigger proinflammatory cytokines was observed with its administration inducing a pro-inflammatory and pro-apoptotic profile, evident from the significant elevation of TNF-α, decrease and caspase-3 compared to control. Elevated TNF-α reflects activation of systemic inflammatory pathways, while increased caspase-3 indicates enhanced apoptotic signaling. Cyclophosphamide administration also significantly decreased IL-10 levels compared to control, indicating suppression of anti-inflammatory signaling. This finding underscores CYP’s strong pro-inflammatory effect, as reduced IL-10 would limit the body’s ability to counterbalance cytokine-driven inflammation. Treatment with MP extract alone at both 25 and 50 mg/kg produced a distinct immunomodulatory pattern with the decrease in TNF-α suggesting suppression of pro-inflammatory cascades, while the IL-10 and Caspase-3 responses are consistent with the upregulation of anti-inflammatory mechanisms and protection against apoptotic cell death respectively. This profile reflects a shift toward an anti-inflammatory and cytoprotective state, aligning with MP’s adaptogenic and neuroprotective properties reported in ethnopharmacology [28]. In the CYP/MP co-treatment groups, Muira puama demonstrated marked protective effects including TNF-α levels being significantly reduced compared to CYP alone, showing attenuation of CYP-induced inflammation, IL-10 levels were further elevated relative to CYP, indicating potentiation of anti-inflammatory signaling and caspase-3 levels were significantly reduced compared to CYP, suggesting mitigation of apoptotic processes. Together, these effects demonstrate that MP extract effectively counteracts CYP-induced inflammatory stress and apoptosis, restoring immune balance by downregulating TNF-α while upregulating IL-10, and protecting cellular integrity through caspase-3 inhibition.

In this study, CYP-treatment resulted in significant suppression of plasma testosterone, luteinizing hormone (LH), and follicle-stimulating hormone (FSH) compared to control. This pattern is consistent with CYP’s known gonadotoxic and endocrine-disrupting properties, which impair the hypothalamic–pituitary–gonadal (HPG) axis and testicular steroidogenesis [2]. Administration of Muira puama (MP25 and MP50) alone significantly increased plasma testosterone levels, suggesting a stimulatory effect on androgen production. This aligns with the traditional use of Muira puama as an aphrodisiac and tonic, and may reflect enhanced testicular steroidogenesis or modulation of gonadotropin signaling [8, 28, 29]. When co-administered with CYP (CYP/MP25 and CYP/MP50), testosterone levels were significantly restored compared to CYP alone, highlighting MP’s protective effect against CYP-induced hypogonadism. For gonadotropins, CYP treatment significantly decreased both LH and FSH compared to control, indicating disruption of pituitary regulation. Co-treatment with MP partially reversed these effects: LH and FSH levels were significantly higher in CYP/MP25 and CYP/MP50 compared to CYP alone, although not fully restored to control levels. This suggests that MP may support pituitary function and testicular feedback loops under conditions of CYP-induced suppression.

Cyclophosphamide treatment also resulted in a reduction in gonadosomatic index compared to control, reflecting gonadal atrophy and impaired reproductive organ development. This aligns with the well-known gonadotoxic potential of CYP, which induces testicular degeneration through oxidative stress, inflammation, and apoptosis of germ cells [2]. The co-administration of Muira puama (CYP/MP25 and CYP/MP50) significantly increased GSI relative to CYP alone, indicating a protective effect. Although GSI values in these groups remained lower than control, the partial restoration suggests that Muira puama mitigates CYP-induced testicular damage. The improvement in GSI correlates with the observed restoration of plasma testosterone, LH, and FSH levels as well as the reduction in oxidative stress and apoptosis. Together, these findings suggest that Muira puama preserves gonadal mass and function by counteracting CYP-induced cellular damage and endocrine suppression.

Histological analysis of the control group revealed normal testicular architecture, including intact seminiferous tubules, orderly spermatogenic cell layers, well-defined basement membranes, and abundant Leydig and Sertoli cells. These features are characteristic of healthy testes with preserved spermatogenesis [30]. Treatment with CYP resulted in marked gonadotoxic alterations, including disorganization of seminiferous tubules, reduced germ cell density, and basement membrane disruption. Such changes reflect CYP-induced oxidative stress, inflammation, and apoptotic injury, consistent with the biochemical findings in this study. In contrast, co-treatment with Muira puama demonstrated structural preservation of the testes. These sections showed better organization of seminiferous tubules, improved Leydig cell morphology, and the presence of active spermatogenic cells. While not fully identical to control, the restoration observed in CYP/MP groups indicates that Muira puama confers partial histological protection against CYP-induced testicular degeneration.

**Plate 1:**
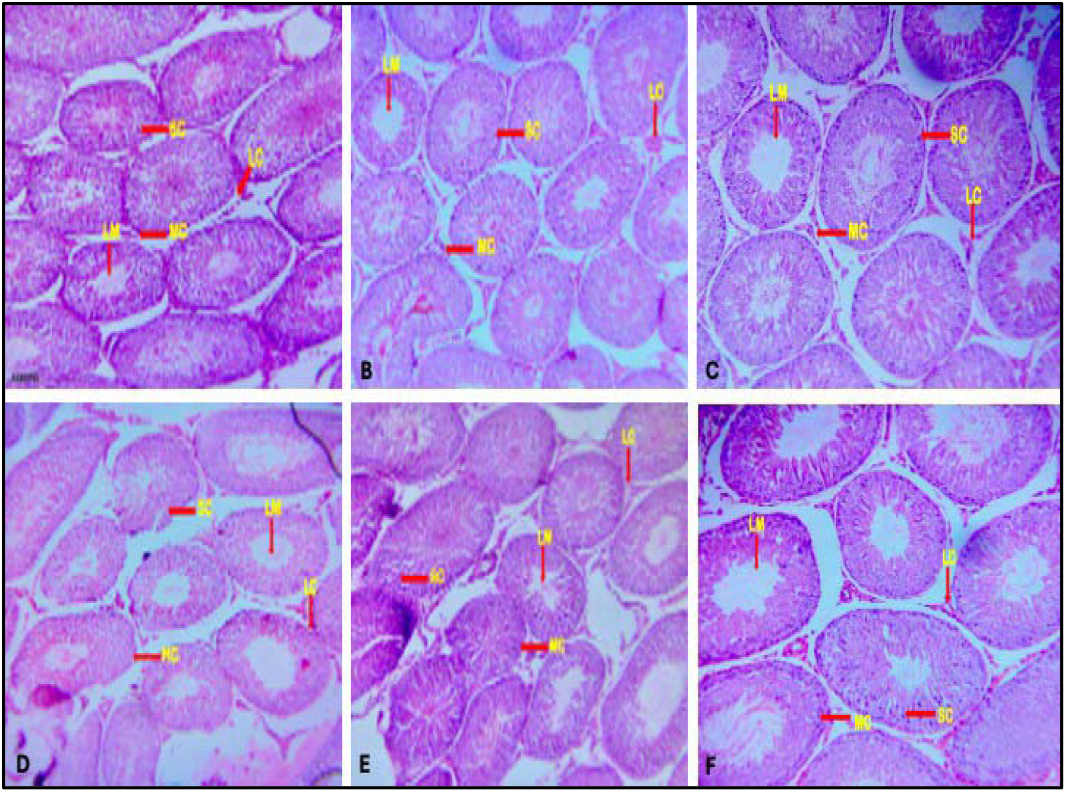
Photomicrograph of testes-stained section by haematoxylin and eosin X100 Stained slides revealed testicular tissue of control (A), MP25 (B), MP50 (C), CYP (D), CYP/MP25 (E), CYP/MP50 (F). MC: Myoid Cell, L: Lumen, LC: Leydig Cell, Sc: Sertoli cell Mag X400.

## Conclusion

This research demonstrated the protective effects of Muira puama extract against cyclophosphamide induced gonadotoxicity in rats. However, additional research is necessary to examine its molecular mechanisms and possible clinical applications.

## Declaration of Ethics approval

Ethical approval was obtained from the Faculty of Basic Medical Sciences LAUTECH

